# Disambiguating glutamatergic and cholinergic signaling under ketamine in *C. elegans*

**DOI:** 10.1101/2025.08.02.667727

**Authors:** Belen Karakullukcu, Hamilton White, Christopher V Gabel, Christopher W Connor

## Abstract

**Background:** Although classed as an N-methyl-D-aspartate (NMDA) receptor antagonist, ketamine acts on other targets with comparable potency, including acetylcholine receptors. Yet, the degree to which these additional actions matter *in vivo* is unknown. We used *Caenorhabditis elegans*, in which the NMDA receptor subunit NMR-1 can be removed genetically, to disambiguate these actions.

**Methods:** Wild-type and *nmr-1* null animals were scored for immobility and insensitivity to touch after 1 hour in 0–50 mg/mL ketamine. Cholinergic neuromuscular transmission at 0–15 mg/mL ketamine was assessed by the rate to paralysis via aldicarb (cholinesterase inhibitor) and immobilization via tetramisole (cholinergic agonist). System-wide neuronal activity was measured via fluorescence calcium imaging across 120 head neurons (0–7.5 mg/mL) and individually in the interneuron AVA (5 mg/mL).

**Results:** Ketamine produced concentration-dependent immobility (wildtype, 30.7±3.4% at 5 mg/mL to 96.0±1.6% at 50 mg/mL); *nmr-1* null animals were more sensitive to ketamine immobility (48.0±3.9% at 5 mg/mL; p=0.012 *versus* wildtype). Ketamine counteracted tetramisole-induced paralysis (median movement 1.13 *versus* 0.29; p<0.001) and delayed aldicarb-induced paralysis in both genotypes (50% paralysis at, wildtype 97.4±4.0 to 150.5±4.2 min, p<0.001; *nmr-1* 111.3±6.6 to 163.1±12.5 min, p=0.034). Multi-neuron imaging showed that ketamine shifted system activity toward more frequent transitions and shorter-lived system states (p=0.029–0.038). AVA, a key interneuron with NMDA receptor expression, also displayed this shift in dynamics in WT animals exposed to ketamine (0.20 to 0.37 events/min, p=0.004; mean duration 106 to 56 s, p=0.001), but not in *nmr-1* null animals (p≥0.93).

**Conclusions:** In *C. elegans* ketamine acts through at least two distinct mechanisms: an NMR-1-dependent change in nervous system state dynamics and a separate action on cholinergic neuromuscular signaling that accompanies immobilization. Thus, multi-target descriptions are required to understand the *in vivo* effects of ketamine even in the simple *C. elegans* model system and should be considered in higher organisms including humans.

**Summary Statement:** In *Caenorhabditis elegans*, ketamine acts through at least two separable mechanisms: an NMDA receptor-dependent change in the timing and periodicity of neuronal activity, and an NMDA receptor-independent action on cholinergic signaling that accompanies immobility.

## Introduction

Since its introduction as a dissociative anesthetic, ketamine’s antagonistic effects on NMDA have been the focus of our pharmacological understanding of its clinical properties.^1–3^ However, its adjunct effects – such as acute opioid-sparing analgesia, treatment of neuropathic pain, or intervention in treatment-resistant depression, which persist for longer than the pharmacokinetic presence of the drug itself – suggest that NMDA antagonism alone is an insufficient explanatory model.^4–6^ Beyond the NMDA receptor, ketamine acts on opioid, nicotinic and muscarinic acetylcholinergic, HCN1, and 5-HT2 receptors, among others.^7,8^ This promiscuity in receptor binding makes the unitary/monogamous language used to describe anesthetic agents misleading. Ketamine is actually most potent at the neuronal α4β4 nicotinic receptor (IC_50_ 0.144μM), rather than at NMDAR (IC_50_ 0.338μM).^8^

*In vitro* potency cannot tell us which receptor interactions are most important in a behaving animal. Instead, specific targets need to be isolated systematically and studied *in vivo*. The nematode *C. elegans* is an established model system for anesthesia research and uniquely suited to addressing these questions, offering a well-characterized nervous system, tractable *in vivo* experiments, and expansive genetic tools.^9–13^ *C. elegans* possess NMDA-type glutamate receptors and a large, completely-mapped nicotinic system. We already know what specific behavioral changes are produced by isolated NMDAR antagonism: the behavioral effects of nitrous oxide (N_2_O) require NMR-1, the GluN1 subunit of NMDAR.^14^ NMDAR antagonism with 70% N_2_O produces only an isolated defect in the reversal response to noxious stimulus. N_2_O does not cause immobility, nor even slow locomotion, and cholinergic transmission is unaltered.^13^ *C. elegans nmr-1* null animals are unaffected by N_2_O.

In *C. elegans,* ketamine 20mM (∼5.5mg/mL) causes significant immobilization without activating a stress response,^15^ and long-term larval exposure produces durable behavioral changes.^16^ Here, we pair multi-neuron imaging in young adult *C. elegans* with behavioral pharmacology. Using *nmr-1* null animals, we eliminate the primary NMDAergic effect and elucidate other actions, employing comprehensive system-wide neuron imaging with single neuron resolution. Additionally, we image the AVA neuron alone, a cholinergic interneuron that receives glutamatergic input via NMDAR and synapses onto motor neurons via acetylcholine.^14,17^ In addition to being an NMDAR antagonist, ketamine alters cholinergic signaling.^4–6,18,19^ We perform ET50 assays to distinguish interactions between ketamine-induced paralysis versus paralysis caused by the cholinesterase inhibitor aldicarb and the acetylcholine agonist tetramisole.

Ultimately, we show two distinct and separable behavioral and neurological actions of ketamine in one animal: an NMDA receptor-dependent neuronal signature and a cholinergic behavioral action.

## Methods

### Strain selection

*C. elegans* were maintained at 20°C on nematode growth medium (NGM) plates seeded with Escherichia coli OP50 using standard methods.^20^ These strains of young adult hermaphrodites in the N2 background were used:

- QW1144: lin-15(n765ts); zfIs124[Prgef-1::GCaMP6s] II
- OH10689: otIs355[Prab-3::NLS::tagRFP] IV
- QW625: zfIs42[Prig-3::GCaMP3::SL2::mCherry]
- VM487: nmr-1(ak4) II

For multi-neuron imaging, QW1144 and OH10689 were crossed to combine pan-neuronal cytoplasmic GCaMP6s with pan-neuronal nuclear tagRFP (QW1144;OH10689). QW625, expressing GCaMP3 and mCherry in AVA, was crossed into VM487; the resulting strain was used for all *nmr-1* null behavioral and imaging experiments.^14,17^ In behavioral and aldicarb assays, QW1144;OH10689 served as wildtype and QW625;VM487 as *nmr-1* null. In AVA imaging, QW625 served as wildtype. Neither QW1144 nor QW625 carry the *lite-1* mutation, thus maintaining wildtype-like sensory function in response to blue light. As invertebrates, *C. elegans* are exempt from IACUC review.

### Ketamine concentration and exposure time (Figure 1)

*C. elegans* were exposed to 0, 5, 10 and 50mg/mL ketamine. For each trial, 15 animals were placed in a well containing 500μL of S-basal (100 mM NaCl, 50mM potassium phosphate buffer, 5μg/mL cholesterol) with ketamine (Covetrus, USA). Ketamine’s behavioral inhibition plateaus after 60 min. Animals not moving spontaneously were scored as immobile. Animals not responding to harsh touch with a worm pick were scored as insensate. Five independent trials of 15 animals each were performed per concentration per genotype (total 75 animals per concentration per genotype).

**Figure 1:**
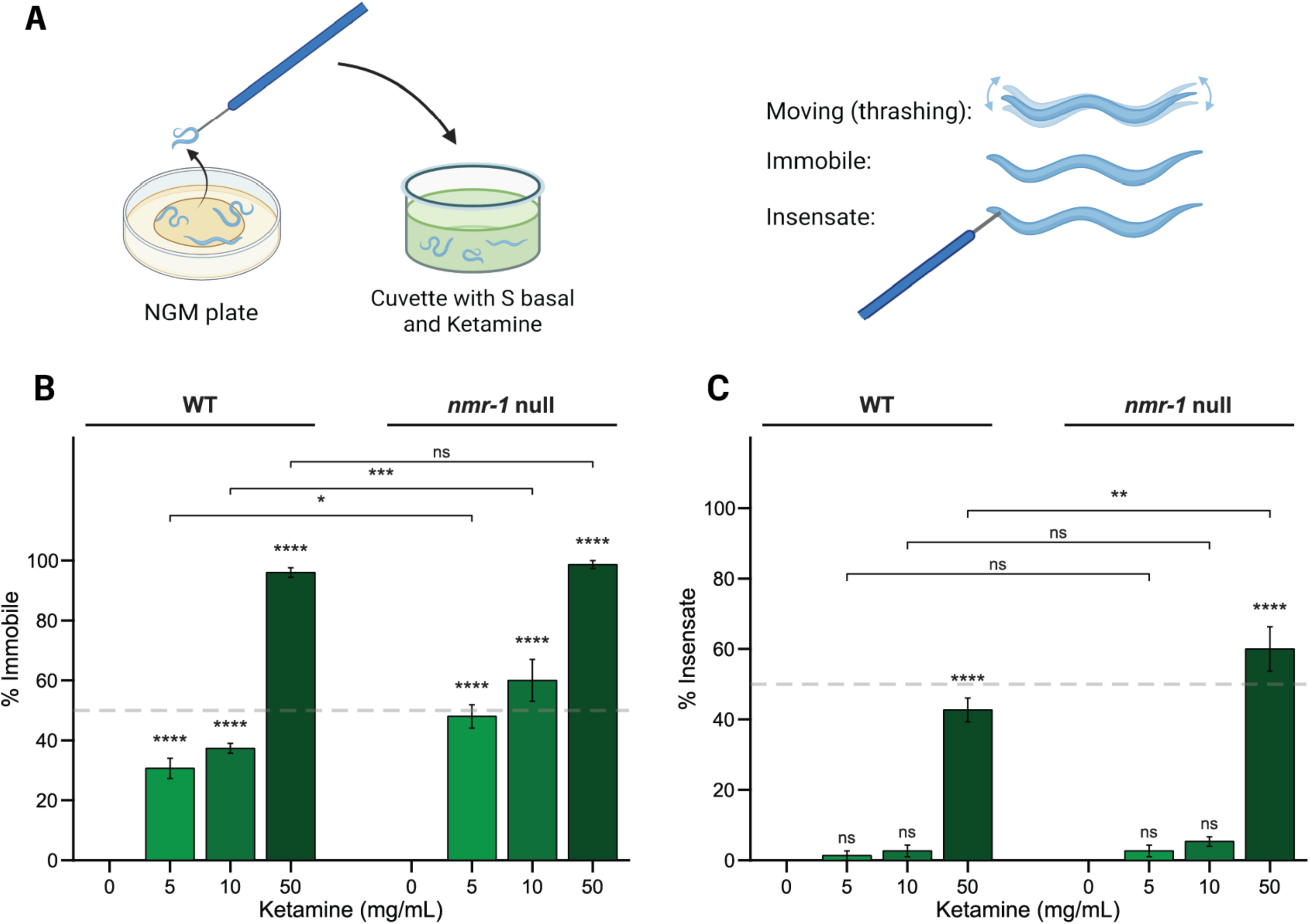
Ketamine-induced immobility and insensitivity. (A) Assay schematic. Adult animals were transferred from NGM plates to a well containing ketamine in S-basal and scored after 1 h as moving (thrashing), immobile (no spontaneous movement but responsive to harsh touch) or insensate (no response to harsh touch). Created in BioRender. (B, C) Percentage of immobile (B) and insensate (C) wild-type and *nmr-1* null animals at 0, 5, 10 and 50 mg/mL ketamine. Bars show mean ± SEM across five independent trials of 15 animals each (75 animals per concentration per genotype); each trial contributes one percentage, so n = 5 trials per bar. Stars above bars: comparison with 0 mg/mL within each genotype (one-way ANOVA followed by Tukey–Kramer). Brackets: comparison between genotypes at the same concentration (two-way ANOVA followed by Tukey–Kramer). Dashed line, 50%. Ns, not significant; * p < 0.05, ** p < 0.01, *** p < 0.001, **** p < 0.0001.

### Aldicarb assay

Aldicarb (1mM) and/or ketamine (0.137–15mg/mL) were added to unseeded NGM plates with standard supplements (MgSO_4_, CaCl_2_, KPO_4_, cholesterol). 15 animals were transferred to each plate and scored every 15 min until all scoreable animals became paralyzed (no response to three touches to the head).^21^ Animals that burrowed into the agar were excluded from the count at that timepoint.

### Light-sheet imaging

Neuronal activity was imaged with a diSPIM microscope (Applied Scientific Instrumentation, USA) and a water-immersion 40×0.8 NA objective (Nikon, USA). Volumetric images of GCaMP and red fluorescence (tagRFP or mCherry) were acquired at 2 Hz with a voxel size of 0.1625×0.1625×1 μm. GCaMP and the red fluorophores were excited at 488 and 561 nm, respectively (10 mW; Vortran Laser Technology, USA). Animals were immobilized in a permeable hydrogel of 13.3% polyethylene glycol diacrylate (Advanced BioMatrix, USA) and 0.1% Irgacure (Sigma-Aldrich, USA) containing tetramisole (Sigma-Aldrich, USA).^22^ Tetramisole at low to moderate concentrations is widely used to immobilize *C. elegans* for neuronal imaging.^23^ Tetramisole 7.5mM (above the 5mM used here) does not significantly alter axonal transport in AWC neurons.^24^ We have published these imaging methods previously.^9–11,25,26^

Hydrogel pads were cured onto silanized coverslips with ultraviolet light, and coverslips were secured with vacuum grease to the bottom of a 50-mm Petri dish filled with 50 mL of immersion medium (S-basal with 5mM tetramisole and ketamine as above).

### Imaging protocols

*Induction (Figure 2).* After 1 hour in hydrogel with 5mM tetramisole, 5mg/mL ketamine was added to the medium and imaging began immediately, continuing for 131 min. This interval captures initial induction over 60 minutes plus the subsequent steady state.

**Figure 2:**
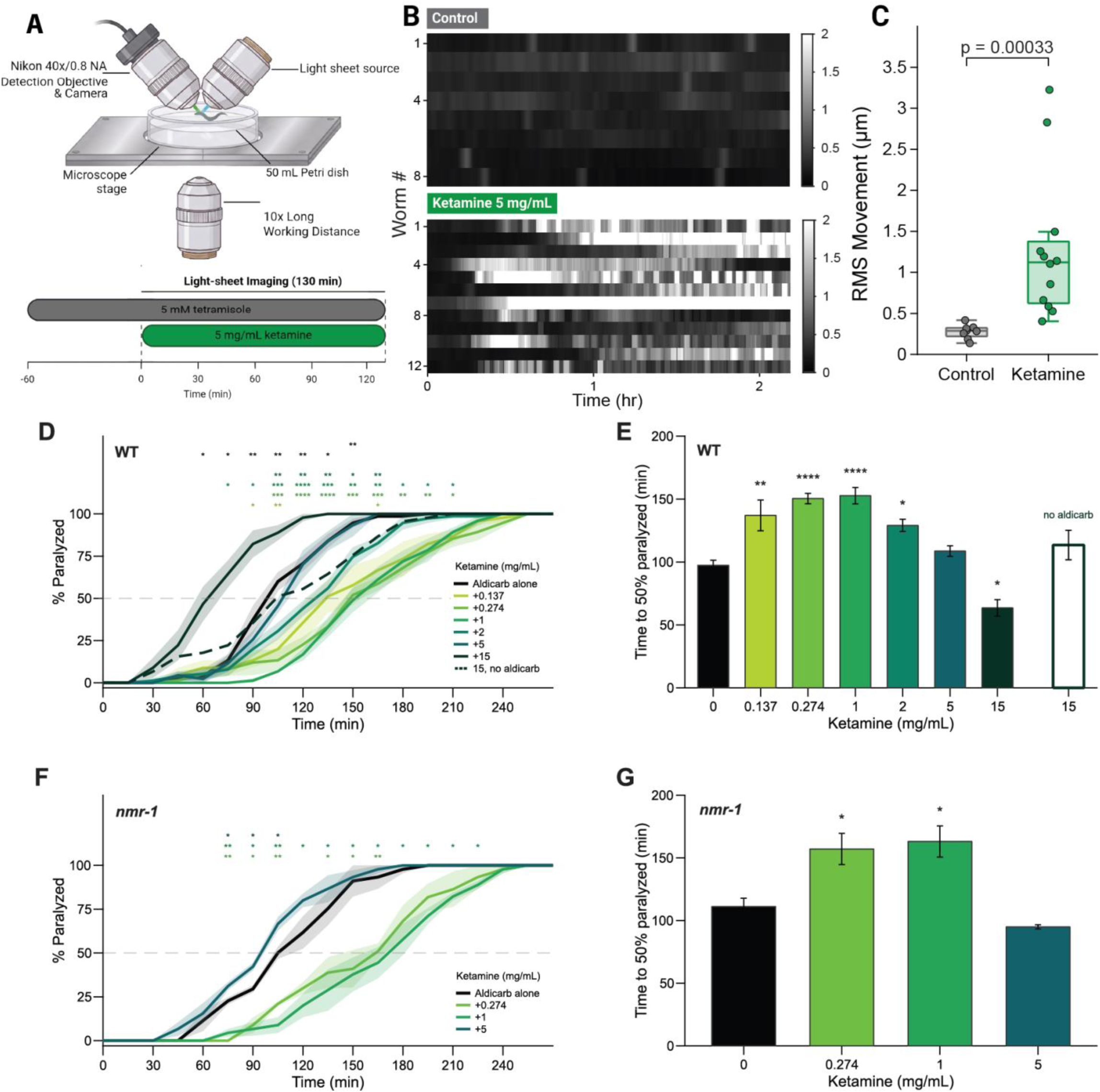
Ketamine alters movement and aldicarb sensitivity. (A) Light-sheet imaging setup for neuronal imaging and protocol. Animals were immobilized in hydrogel with 5 mM tetramisole for 60 min, after which 5 mg/mL ketamine was added, and animals were imaged for 130 min. Created in BioRender. (B) RMS movement of each animal over the recording (2-min sliding window) for control (top, n=8) and ketamine-treated (bottom, n=12) animals. Each row is one animal; color scale, RMS movement in μm (C) Mean RMS movement (um) of each animal from 60 min after the start of recording to the end. Boxplots show the median and interquartile range, with whiskers extending to the most extreme values within 1.5 × the interquartile range; points are individual animals. Wilcoxon rank-sum test. (D, F) Percentage of paralyzed wild-type (D) and *nmr-1* null (F) animals over time on 1 mM aldicarb alone (black) or with ketamine (green; concentrations in mg/mL). Lines and shading show mean ± SEM across trials. The dashed dark green line in (D) is 15 mg/mL ketamine without aldicarb, a ketamine-alone control (n=3 trials), shown without shading for clarity. Stars, color-coded by ketamine concentration, show comparisons with aldicarb alone at each timepoint (Welch’s t-test). Dashed gray line, 50% paralyzed. (E, G) Time to 50% paralysis (ET50) for wild-type (E) and *nmr-1* null (G) animals, estimated for each trial by linear interpolation. Bars show mean ± SEM; stars show comparisons with aldicarb alone (one-way ANOVA followed by Tukey-Kramer). The open bar in (E), set apart on the right, is the ketamine-alone control (15 mg/mL, no aldicarb). Every trial plated 15 scoreable animals and contributes one ET50, so n is the number of trials: wild type, n=5 trials for aldicarb alone, 0.274, 1 and 2 mg/mL and n=3 trials for 0.137, 5 and 15 mg/mL and for ketamine alone; nmr-1, n=3 trials per condition. * p < 0.05, ** p < 0.01, *** p < 0.001, **** p < 0.0001.

*Steady state (Figure 3).* Animals were incubated in the imaging medium and 0, 2.5, 5 or 7.5mg/mL ketamine for 1 hour and then imaged for 11 min. The first 2 min of each recording were excluded from analysis.

**Figure 3:**
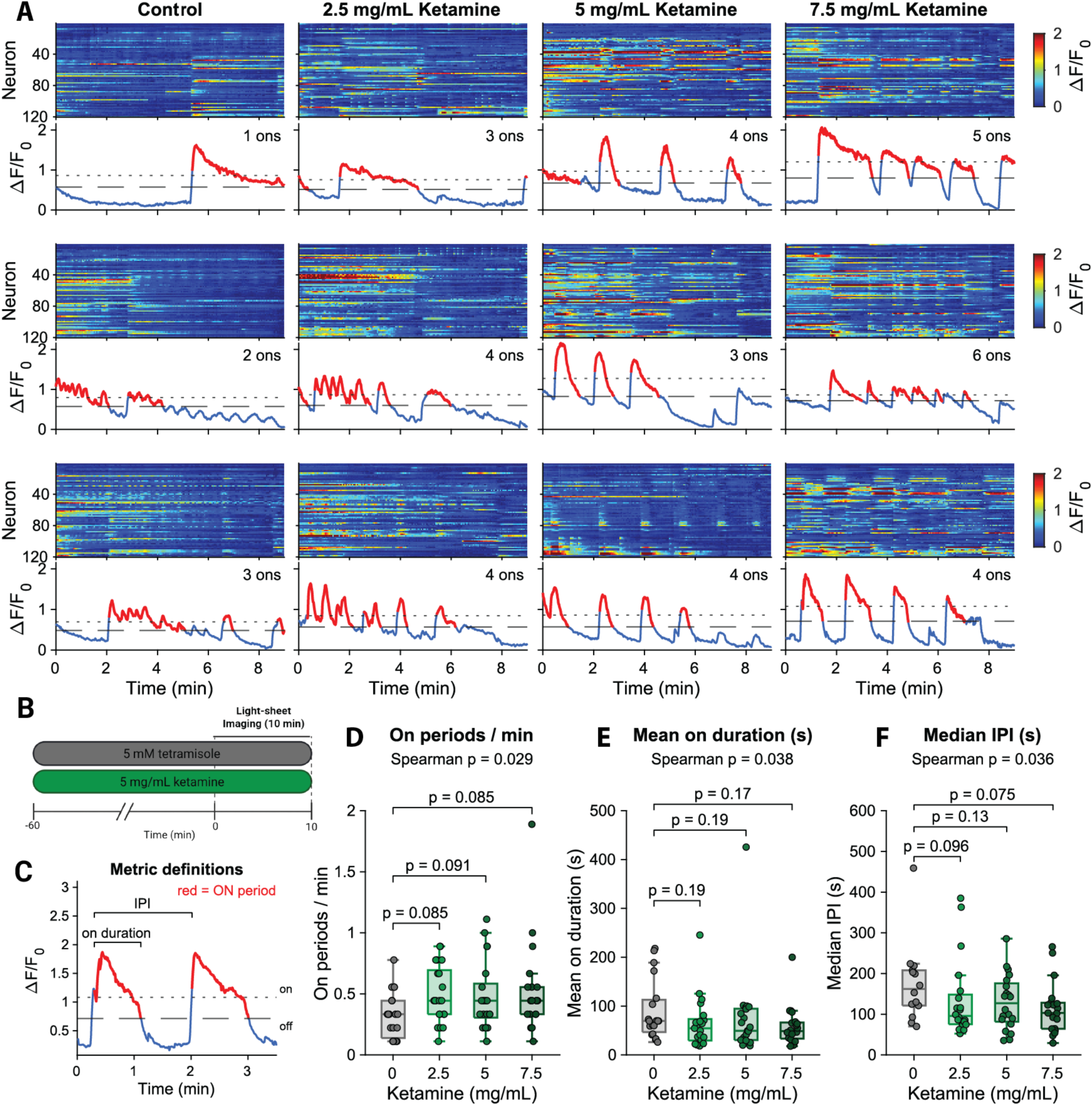
Patterns of neuronal activity in response to ketamine exposure. (A) Three example recordings at each ketamine concentration. For each recording, the heatmap shows ΔF/F0 of the 120 tracked neurons, and the trace below shows the network activity trace (mean activity of the largest group of correlated neurons) with ON periods in red and OFF periods in blue. Dotted and dashed lines mark the ON and OFF thresholds; the number of ON periods is shown at the top right. Examples shown: control, W5, W19 and W13; 2.5 mg/mL, W1, W4 and W12; 5 mg/mL, W11, W21 and W3; 7.5 mg/mL, W5, W4 and W1. (B) Protocol. Animals were incubated with 5 mM tetramisole and ketamine (0, 2.5, 5 or 7.5 mg/mL) for 60 min and imaged for 10 min; the first 2 min were excluded from analysis. Created in BioRender. (C) Metric definitions, shown on the first two ON periods of a 7.5 mg/mL recording (W1). ON periods were defined with a hysteretic threshold (see methods) ON duration, time from an ON onset to the following OFF; IPI (inter-ON-period interval), time between consecutive ON onsets. (D–F) ON periods per minute (D), mean ON-period duration (E) and median IPI (F) for each recording. Box plots as in Figure 2C; points are individual recordings. Spearman p values (top) are from one-sided Spearman rank correlations between ketamine concentration and each metric, with the direction specified a priori. Brackets show comparisons with control (Wilcoxon rank-sum, Holm-corrected). n = 19, 21, 21 and 20 recordings at 0, 2.5, 5 and 7.5 mg/mL; for median IPI, n = 14, 20, 20 and 19 recordings with at least two ON periods.

*AVA (Figure 4).* Animals were incubated in 5mM tetramisole and 0 or 5mg/mL ketamine for 1 hour and then imaged for 31 min; the first minute of each recording was excluded for photobleaching. The 30-min recording window allows multiple AVA ON/OFF transitions to be captured.^27^

**Figure 4:**
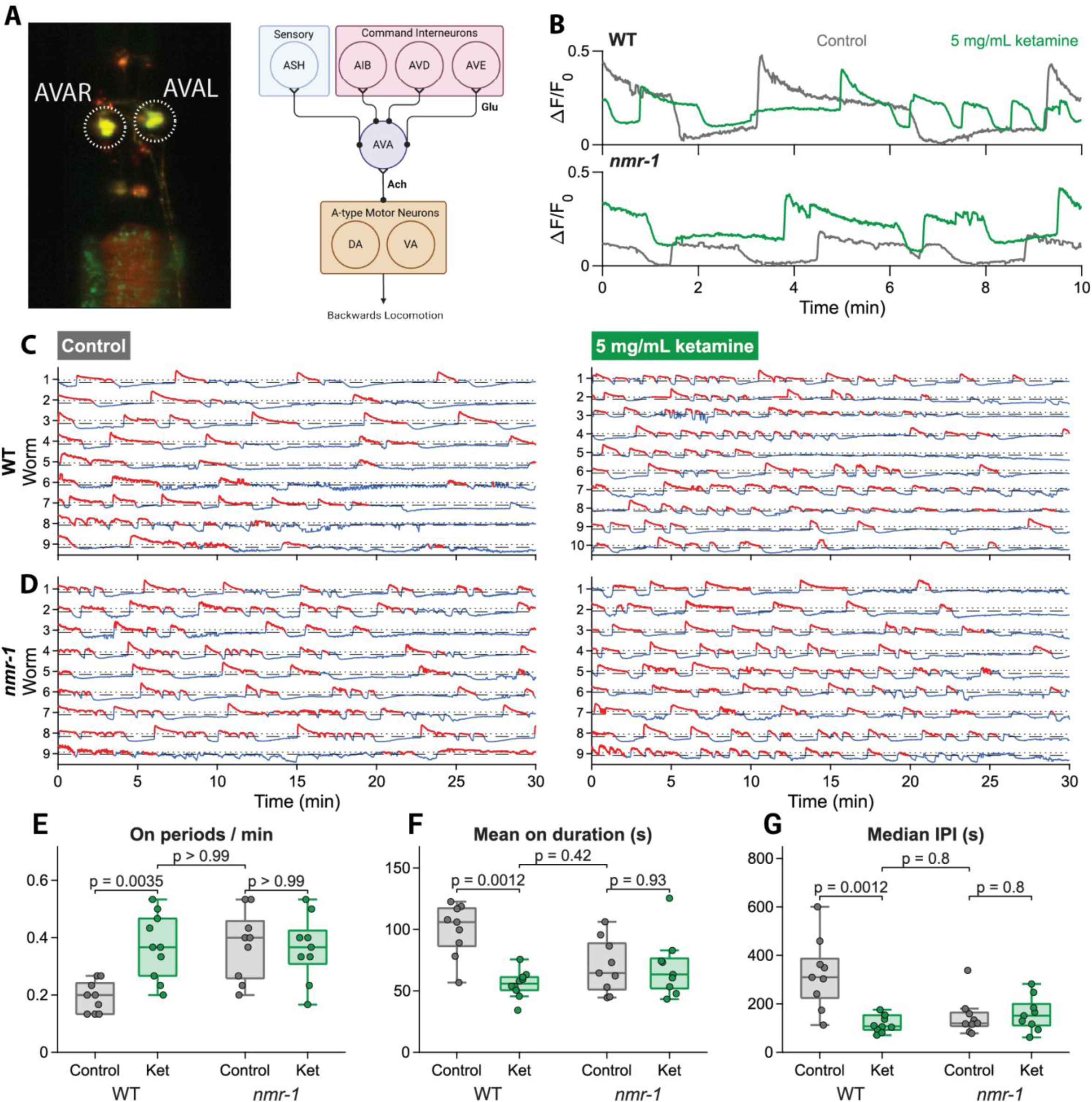
Ketamine alters AVA activity through NMR-1. (A) At left, the AVA neuron pair (AVAL and AVAR, dotted circles) as imaged in an example animal. At right, simplified circuit: AVA receives glutamatergic input from sensory neurons and command interneurons and makes both cholinergic synapses and gap junctions onto A-type motor neurons, driving backward locomotion events. Created in BioRender. (B) Example AVA activity (mean ΔF/F0 of the AVAL and AVAR pair) in control (gray) and 5 mg/mL ketamine-treated (green) wild-type (top) and *nmr-1* null (bottom) animals. The first 10 min of each recording are shown; traces were normalized over the full recording. (C, D) Collective AVA activity over the full 30-min recording for every imaged wild-type (C) and *nmr-1* null (D) animal, control (left) and ketamine-treated (right), with ON periods in red and OFF periods in blue. Dotted and dashed lines mark the ON and OFF thresholds. (E–G) ON periods per min (E), mean ON-period duration (F) and median IPI (G). Boxplots as in Figure 2C; points are individual animals. Brackets show Wilcoxon rank-sum tests between control and ketamine within each genotype, and between ketamine-treated wildtype and untreated *nmr-1* null animals. Holm-corrected across the three comparisons within each metric. Wild type, n = 9 control and 10 ketamine; *nmr-1*, n = 9 per condition.

## STATISTICAL METHODS

### Extraction of neuronal activity

For each QW1144;OH10689 animal imaged, 120 neurons were tracked in the head region using nuclear-localized RFP with their activity measured by cytoplasmic GCaMP6s fluorescence in the immediately surrounding volume. Similarly, in QW625 and QW625;VM487, the activity of the left and right AVA neurons (AVAL, AVAR) were extracted. Tracking and extraction were performed using our published automated pipeline.^11,26^

### Immobility and insensitivity (Figure 1)

The percentage of immobile and insensate animals at each ketamine concentration was calculated relative to the number of animals in the well, shown as mean±SEM. Each trial contributes one percentage; trials were treated as independent replicates (n=5 trials/condition). Within and across genotypes, the effect of concentration was tested with one-way or two-way ANOVA, respectively.

### Movement during whole-brain imaging (Figure 2B–C)

Movement was calculated from the tracked 3D positions of 120 neurons in each recording. At each timepoint, the interval distance moved by each neuron was averaged across neurons and converted to a root-mean-square (RMS) movement over a sliding 2-min window. At stable state, each animal’s RMS movement was averaged from 60 min after the start of recording to the end (∼last 70 min) and compared with a two-sided Wilcoxon rank-sum test.

### Aldicarb paralysis (Figure 2D–G)

At each timepoint, % paralyzed was calculated as the numerator of paralyzed animals divided by the denominator that could be scored, excluding animals burrowed into the agar. Trials of 15 animals were followed until every scoreable animal was paralyzed. Time courses show mean±SEM % paralyzed across trials; at each timepoint, each ketamine concentration was compared with aldicarb alone using a two-sided Welch’s t-test. For each trial, the time to 50% paralysis (ET50) was estimated is shown as mean±SEM across trials. Within each genotype, ET50 values were compared across conditions by one-way ANOVA and Tukey–Kramer. Genotypes were compared by two-way ANOVA (genotype×ketamine concentration, with interaction), followed by Tukey–Kramer.

### Detection of ON/OFF activity states (Figures 3,4 and Supplemental Figure 1,2)

For each recording, the activity trace (mean activity of largest neuron group or AVA, ΔF/F0) was smoothed with a 5-s moving average. ON and OFF thresholds for hysteresis thresholding were set using the 10^th^ and 90^th^ percentiles of the smoothed trace. OFF periods shorter than 5 seconds were merged into the surrounding ON period. Four metrics were calculated for each recording: ON periods per minute, mean ON-period duration, median time between consecutive ON periods (inter-period interval, IPI), and the percentage of time spent in ON periods (duty cycle).

### Whole-brain activity across ketamine concentrations (Figure 3)

The first 2 min of each recording were removed to eliminate initial fluorophore photobleaching. The remaining 9 min were analyzed. Each neuron’s trace was normalized as ΔF/F0 (F0: the mean of the lowest 1% of values). To obtain a single network activity trace per recording, the 40 most active neurons (upper tercile) with the highest standard deviation were selected. Pearson correlations of the time derivative of their ΔF/F0 traces were computed between every neuron pair.^28^ Neurons with correlations greater than 0.6 were considered connected, and the largest group of connected neurons was identified. The mean activity of the neurons within this group was analyzed to measure network dynamics. Initially, 21 recordings were obtained for each concentration. Recordings were excluded if no large connected group was identified, producing n=19, 21, 21 and 20 recordings at 0, 2.5, 5 and 7.5mg/mL respectively. Concentration dependence of group dynamics was tested with a one-sided Spearman rank correlation between ketamine concentration and the metric, with the direction of each test chosen *a priori* from the expected effect of ketamine (ON periods per minute increasing; ON-period duration and IPI decreasing with concentration). Intergroup differences were tested with Kruskal-Wallis. Each concentration was compared with control using a two-sided Wilcoxon rank-sum test and Holm multiple comparisons correction.

### AVA activity in wild-type and *nmr-1* animals (Figure 4)

Each 31-min recording was analyzed from minute 1 onward (30 min). C. elegans contains a symmetric pair of command interneurons, AVAL and AVAR, whose activity is closely correlated. AVAL and AVAR traces were normalized as ΔF/F0. Their mean signal was used as the AVA trace for that animal. Control and ketamine-treated recordings were compared within each genotype, and ketamine-treated wildtype was compared with untreated nmr-1, using two-sided Wilcoxon rank-sum tests and Holm multiple comparisons correction.

### AVA activity in microbead-immobilized animals (Supplemental Figure 1)

Animals were immobilized on a 5% agarose pad under a coverslip with 5μL of a 0.65% (w/v) suspension of 0.051-μm polystyrene microspheres, without tetramisole.^29^ The fluorescence of one AVA neuron (AVAL or AVAR) per animal was measured in each frame and normalized to ΔF/F0. ON/OFF periods were detected as above, and control and ketamine-treated recordings were compared with two-sided Wilcoxon rank-sum tests.

### Presentation and significance

Boxplots show the median and interquartile range (IQR), with whiskers extending to the most extreme values within 1.5×IQR; points are individual recordings or animals. Bar graphs show mean±SEM. p<0.05 was considered significant. Exact p-values are reported where space allows; otherwise * p<0.05, ** p<0.01, *** p<0.001 and **** p<0.0001.

## Results

### Behavioral effects of ketamine

Ketamine decreases mobility in wild-type N2 *C. elegans*.^15^ We reproduced this effect in our fluorescent imaging strains, and also in *nmr-1* null genotype crosses (Figure 1A). Without ketamine, no animals were immobile or insensate (Figure 1B-C). Ketamine immobilized wild-type animals in a concentration-dependent manner (Figure 1B; p=4.4x10^-15^). 30.7±3.4% of animals were immobile at 5mg/mL (p=7.2x10^-8^), 37.3±1.6% at 10 mg/mL (p=4.2x10^-9^) and 96.0±1.6% at 50mg/mL (p=6.5x10^-18^). *nmr-1* null animals were also immobilized in a concentration-dependent manner (Figure 1B; p=1.4x10^-10^): 48.0±3.9% at 5mg/mL (p=1.7x10^-6^), 60.0±7.0% at 10mg/mL (p=8.1x10^-8^) and 98.7±1.3% at 50mg/mL (p=4.7x10^-11^). Immobility differed between genotypes (genotype p=4.8x10^-5^, concentration p=1.0x10^-23^, interaction p=0.0025). *nmr-1* null animals were more immobilized than wild-type at 5mg/mL (p=0.012) and 10mg/mL (p=0.00050), but at 50mg/mL (p=0.999) both genotypes were maximally immobilized. Insensitivity to noxious stimulus occurred only at the highest concentration (Figure 1C). At 50mg/mL, 42.7±3.4% wild-type (p=3.8x10^-10^) and 60.0±6.3% *nmr-1* null animals (p=4.8x10^-9^) were insensate. Insensitivity differed between genotypes only at 50mg/mL, where more *nmr-1* null than wild-type animals were insensate (p=0.0022, Figure 1C).

### Ketamine and cholinergic signaling

The micro-scale movement of *C. elegans* immobilized by a combination of hydrogel and tetramisole was measured by tracking neuron position using our established neuronal imaging platform (Figure 2A). Whereas movement in control animals remained low throughout the recording, animal movement increased over the first 30 min of exposure to ketamine and remained elevated thereafter (Figure 2B). Median RMS movement was elevated in ketamine-treated animals compared with controls (p=0.00033; Figure 2C).

To probe cholinergic signaling in ketamine-induced immobilization further, we measured sensitivity to aldicarb, a cholinesterase inhibitor that paralyzes animals through acetylcholine accumulation at the neuromuscular junction. Time to paralysis of *C. elegans* placed on aldicarb is a well-established metric of acetylcholine signaling: animals with reduced cholinergic signaling display delayed paralysis.^21^ In wild-type animals, ketamine delayed paralysis at low and intermediate concentrations, shifting the paralysis curves to later times (Figure 2D). The time to 50% paralysis (ET50) increased from 97.4±4.0 min with aldicarb alone to 137.1±12.2 min with 0.137mg/mL ketamine (p=0.0070), 150.5±4.2 min with 0.274mg/mL (p=2.8x10) and 152.8±6.5 min with 1mg/mL (p=1.5x10). At higher concentrations, this delay was progressively lost: ET50 fell to 129.1±4.9 min with 2mg/mL (p=0.015) and to 108.7±4.2 min with 5mg/mL, which no longer differed from aldicarb alone (p=0.93). 15mg/mL accelerated paralysis to 63.5±6.6 min (p=0.029) (Figure 2E). The loss of the delay was itself significant: ET50 at 5mg/mL was 41.8 min shorter than at 0.274mg/mL (p=0.0041) and 44.1 min shorter than at 1mg/mL (p=0.0023). Ketamine alone at 15mg/mL also paralyzed animals, with an ET50 of 113.5±11.7 min, not significantly different from 1mM aldicarb alone (p=0.69; Figure 2E). *nmr-1* null animals showed the same pattern (Figure 2F,G). ET50 increased from 111.3±6.6 min with aldicarb alone to 157.2±12.5 min with 0.274mg/mL ketamine (p=0.036) and 163.1±12.5 min with 1mg/mL (p=0.020), and the delay was again lost at 5mg/mL (95.0±1.7 min, p=0.63). Comparing WT and nmr-1, ketamine strongly affected both genotypes, but we detected no effect from genotype (p=0.39) and no genotype×ketamine interaction (p=0.26). ET50 for wildtype and *nmr-1* animals did not differ at any matched concentration (p≥0.83).

### Comprehensive multi-neuron imaging of ketamine’s effect on global network dynamics

Simultaneously imaging 120 individual neurons in the head of *C. elegans* (out of ∼180 total), we observe characteristic system-state transitions in which large groups of neurons are activated or suppressed in a coordinated manner in both control and ketamine exposed animals (Figure 3A). These dynamics were quantified by identifying the largest group of strongly correlated neurons (r≥0.6) and averaging their activity into a summative network activity trace over 9 minutes (Figure 3B). Each timepoint was classified as ON or OFF for that global state (Figure 3A,C). Overall, ketamine produced a greater number of ON periods/min (p=0.029), shorter ON periods (p=0.038), and decreased interval between ON periods (p=0.036), all in a concentration-dependent manner (Figure 3D-F). However, duty cycle (i.e. the percentage of time in the ON state) did not vary with ketamine concentration (p=0.40; Supplemental Figure 2A).

### NMR-1 and ketamine’s effect on AVA activity

The central command interneuron AVA receives glutamatergic input from sensory neurons and other command interneurons and is known to drive backward locomotion.^14,30–32^ Imaging animals with AVA-specific GCaMP expression allowed us to quantify major system state transitions (Figure 4A).^14^ The AVA pair (AVAL and AVAR) were imaged in wild-type and *nmr-1* null animals for 30 min with or without 5mg/mL ketamine (examples in Figure 4B) and segmented into ON/OFF periods (Figure 4C,D). In wild-type animals, ketamine modified AVA activity in the same direction as the global system network. AVA switched ON more often (p=0.0035; Figure 4E), ON periods became shorter (p=0.0012; Figure 4F), and the median IPI decreased (p=0.0012; Figure 4G). In *nmr-1* null animals, no differences were measured with and without ketamine exposure (Figure 4E–G). However, untreated *nmr-1* null animals demonstrated similar AVA activity to ketamine-treated wildtype: none of three metrics differed between untreated *nmr-1* null and ketamine-treated wild-type animals (Figure 4E–G). AVA duty cycle did not differ with ketamine in either genotype, nor between ketamine-treated wild-type and untreated *nmr-1* nulls (all p≥0.47; Supplemental Figure 2B).

Finally, isolated imaging of the AVA neurons alone allowed for imaging without the use of tetramisole for immobilization (see Methods), but only physical immobilization in agarose. The effect of ketamine on AVA activity did not depend on the immobilization method used: AVA still switched ON more often (p=0.038; Supplemental Figure 1C) and ON periods were shorter (p=0.027; Supplemental Figure 1D). The median IPI showed a similar trend (not significant; Supplemental Figure 1E). Duty cycle likewise did not differ (p=0.82, Supplemental Figure 2C).

## Discussion

*In vitro*, ketamine binds a long list of receptors, and by potency alone it is as much a nicotinic drug as an NMDAergic one.^4,8^ Here we employed *C. elegans* to disambiguate the effects of these two distinct neurological targets. An NMR-1-dependent action reshapes the timing of system-wide neuronal activity and specifically the command interneuron AVA. A separate, NMR-1-independent action on cholinergic signaling accompanies immobilization and opposes aldicarb-induced paralysis.

Ketamine immobilizes *C. elegans* in a concentration-dependent manner. At the highest concentration it abolishes the response to harsh touch (Figure 1). These findings are in agreement with previous *C. elegans* studies.^15^ NMR-1 mutation did not protect animals from ketamine immobility: indeed*, nmr-1* nulls were more sensitive (i.e. immobile) at intermediate concentrations. Here, ketamine is distinct from nitrous oxide, which has an NMR-1-specific effect reducing spontaneous reversal rates in *C. elegans*. Notably, NMDA receptor antagonism by nitrous oxide does not produce immobility in *C. elegans*, nor does it alter the rate of locomotion, chemotaxis, feeding or aldicarb sensitivity.^13^ Ketamine may share an NMR-1-dependent component with nitrous oxide, but its ability to immobilize *C. elegans* must arise independently.

Our results indicate that ketamine-induced immobilization is cholinergic. Two results point to the neuromuscular junction. Firstly, prolonged ketamine treatment reverses the paralytic effects of tetramisole, a widely-used paralytic that hyperactivates the levamisole-sensitive nicotinic receptors of body-wall muscle (Figure 2B, C).^33,34^ Secondly, ketamine delayed aldicarb-induced paralysis at 0.137-2mg/mL in wildtype and at 0.274 and 1mg/mL in *nmr-1* nulls (Figure 2D-G). Increased resistance to aldicarb, a cholinesterase inhibitor, indicates reduced cholinergic transmission.^21^ This aldicarb resistance was unaltered in the *nmr-1* null background, implicating an NMDA-independent mechanism. The differences in ketamine sensitivities in the crawling (aldicarb) vs swimming (tetramisole) assays likely result from experimental factors such as continual ingestion during crawling^35^ and the differing kinematics and underlying neuromuscular activity of the two movements.^36^ Nevertheless, drawing comparisons only within each assay, blockade of nicotinic receptors explains both results. Ketamine blocks mammalian muscle nicotinic receptors by interacting with the open and closed states.^6,37^ It also inhibits neuronal nicotinic receptors at anesthetic concentrations,^5^ and is most potent at the neuronal α4β4 receptor.^8,38^ Inhibition is enhanced once the channel has been opened by agonist,^38^ so an agonist-driven paralysis such as that produced by tetramisole is the condition under which ketamine’s block should be most effective. Nematode muscle receptors contain different subunits,^34^ so we cannot assume the same pharmacology. However, our data show that ketamine’s nicotinic interactions are not a curiosity of *in vitro* binding; they play a key role in the behavior of the whole animal.

Notably, the aldicarb dose–response was biphasic: the ketamine-induced resistance was lost at 5mg/mL and reversed at 15mg/mL, a concentration at which ketamine alone paralyzed animals about as quickly as aldicarb (ET_50_ 114 vs 97 min). The most direct pharmacodynamic interpretation is of two overlapping actions: an anticholinergic effect that first opposes aldicarb-induced paralysis, then ketamine’s own immobilizing effect that causes paralysis. Similar biphasic interactions between ketamine and acetylcholinesterase inhibition (via physostigmine) have been documented in rats,^19,39^ and ketamine inhibits human ganglionic nicotinic receptors stereoselectively at concentrations reached during clinical anesthesia.^40^ In rodents, neuronal nicotinic antagonists enhance ketamine’s analgesia but do not change the dose required for immobility or loss of righting.^41^ Clinically, ketamine’s effects are usually attributed to NMDA receptor antagonism, but our data in *nmr*-1 null *C. elegans* indicate that cholinergic effects occur independently of NMDAR signaling.

Ketamine-induced changes in nervous system dynamics are dependent on NMR-1. Ketamine increases the frequency of global system state transitions, resulting in shorter state durations. Ensemble measurements across the majority of head neurons show a modest but consistently graded effect with concentration (Figure 3). Individual measurement of the AVA interneuron, which mediates behavioral state transitions, shows robust shifts toward shorter, more frequent ON states (Figure 4). AVA is a cholinergic interneuron that expresses the glutamate receptors GLR-1 and NMR-1, synapses onto motor neurons, and plays a key role in controlling behavioral crawling states, with AVA activation corresponding to backward locomotion events.^17,30,31^ Ketamine had no measurable effect on AVA dynamics in *nmr-1* null animals. Moreover, untreated *nmr-1* null animals display AVA dynamics that closely resemble ketamine-treated wildtype. These results are consistent with ketamine’s action as a noncompetitive NMDA receptor antagonist, acting in part by open-channel block that shortens NMDA channel mean open time.^3,42^ Brockie *et al*. electrically recorded from AVA directly and found that NMR-1 provides slow, long-lived NMDA currents that are blocked by the NMDA receptor antagonist MK-801. They proposed that these NMDA currents allow sensory inputs arriving within a limited time window to sum within AVA in order to trigger a reversal event.^14^ Loss of this long-lived depolarizing current, whether by mutation or by ketamine, would be expected to shorten AVA’s active states, consistent with the shorter ON periods we observed in both conditions. However, both *nmr-1* null mutation and N_2_O treatment decrease the rate of reversal events of *C. elegans* (i.e., the number of reversal events/min),^13,14^ contrasting with the increased frequency of AVA ON states that we measure with ketamine and *nmr-1*. This discrepancy suggests that loss of the long-lived NMDA current reduces AVA’s capacity to integrate synaptic inputs and commit to a full reversal event in the downstream motor neurons.^14,27^ Thus, even within the *C. elegans* nervous system, ketamine’s action as an NMDA antagonist appears to contribute to a dissociation between the activity of central interneurons and the behavioral states they normally control.

Ketamine stands out among anesthetics because it does not “turn the brain down”. In humans, ketamine anesthesia modifies the interplay between excitation and inhibition to produce a distinctive EEG pattern in which gamma oscillations alternate with slow-delta oscillations.^1,2^ Modeling ketamine at the whole-brain scale, a shortening of excitatory synaptic currents, together with increased neuronal adaptation, is enough to switch simulated brain activity from an asynchronous awake regime into alternating UP and DOWN states.^43^ Our results point to the same principle in a single identified neuron. Removing the slow NMDA current from AVA, by ketamine or by mutation, did not silence it; it broke its activity into shorter, more frequent active states with no detectable change in overall activity (i.e. duty cycle, Supplemental Figure 2). These effects were also reflected in the global state dynamics of the *C. elegans* nervous system. While the scales are vastly different, in both cases ketamine’s NMDA receptor action restructures neuronal activity rather than abolishing it. This sets ketamine apart from isoflurane which, in *C. elegans*, breaks down correlations between neurons and collapses organized global states.^9,11,25^

In *C. elegans*, we find that ketamine reshapes the timing of command interneuron dynamics via the NMDA receptor, and that behavioral slowing is governed primarily by ketamine’s action on cholinergic signaling. Describing ketamine as an NMDAergic anesthetic captures only one of its actions *in vivo*, just as Kale *et al*. argued from *in vitro* binding that single-target labels misrepresent anesthetic pharmacology.^8^ Our results suggest the importance of cholinergic mechanisms in assessing and understanding the behavioral effects of ketamine beyond anesthesia. Ketamine is increasingly used at subanesthetic doses, as an opioid-sparing analgesic and as a rapid-acting treatment for depression. These uses depend on separating these therapeutic actions of ketamine from its dissociative, psychotomimetic and other effects.^7^ As illustrated by the separate behavioral and neuronal actions of ketamine in *C. elegans*, identifying which target drives which desired effect becomes paramount. *C. elegans* gives us a way to pull these actions apart.

## Acknowledgements

*C. elegans* strains QW1144, OH10689 and QW625 were gifts from the Alkema laboratory (UMass, Worcester, MA). VM487 was obtained from the Caenorhabditis Genetics Center (University of Minnesota).

## Supplemental Figure Legends

**Figure S1: Ketamine alters AVA activity in animals immobilized without tetramisole.** Approximately five QW625 animals were placed on a 5% agarose pad with or without 5 mg/mL ketamine and immobilized under a coverslip with 5 μL of a polystyrene microbead suspension (30 μL of 2.6% 0.051-μm beads in 90 μL S-basal; Polysciences cat. no. 08691), as described Kim et al., 2013.^29^ One AVA neuron (AVAL or AVAR) per animal was imaged for 10 min on a Nikon Ti-U inverted fluorescence microscope (1.4 Hz). Fluorescence of AVA and a nearby background region was measured in ImageJ, background-subtracted, and expressed as ΔF/F0, with F0 the mean of the lowest 1% of values.

(A, B) Background-subtracted ΔF/F0 of AVA in every control (A, n = 9) and 5 mg/mL ketamine-treated (B, n = 8) animal, with ON periods in red and OFF periods in blue. Dotted and dashed lines mark the ON and OFF thresholds.

(C–E) ON periods per minute (C), mean ON-period duration (D) and median IPI (E). Box plots as in Figure 2C; points are individual animals. Wilcoxon rank-sum test; a single comparison per metric, so no correction applies.

**Figure S2: Ketamine alters the timing of ON/OFF states without a detectable change in duty cycle.** Duty cycle was calculated for every recording analyzed in Figures 3 and 4 and in Supplemental Figure 1, as the summed duration of all detected ON periods divided by the recording duration, expressed as a percentage. ON/OFF detection and settings were identical to those figures. Box plots as in Figure 2C; points are individual recordings. All three panels share one y-axis. The metrics in Figures 3D–F and 4E–G change substantially in these same recordings, whereas duty cycle does not.

(A) Multi-neuron imaging (from Fig 3): network activity at 0, 2.5, 5 and 7.5 mg/mL ketamine (n = 19, 21, 21 and 20 recordings). Two-sided Spearman rank correlation between ketamine concentration and duty cycle; duty cycle has no a priori direction, so the test is two-sided. No concentration differed from control (Wilcoxon rank-sum, Holm corrected, all p≥0.64).

(B) AVA in wild-type and *nmr-1* null animals, control and 5 mg/mL ketamine (wild type, n = 9 control and 10 ketamine; *nmr-1*, n = 9 per condition). Data from Figure 4. Brackets show Wilcoxon rank-sum tests between control and ketamine within each genotype, and between ketamine-treated wild type and untreated *nmr-1* null animals, Holm-corrected across the three comparisons, as in Figure 4E–G.

(C) AVA in microbead-immobilized animals, control (n=9) and 5 mg/mL ketamine (n=8). Data from Figure S1. Wilcoxon rank-sum test.

