## Supplementary figures and images for "Disambiguating glutamatergic and cholinergic signaling under ketamine in *C. elegans*"

### Supplemental Figure 1

**Supplemental Figure 1**

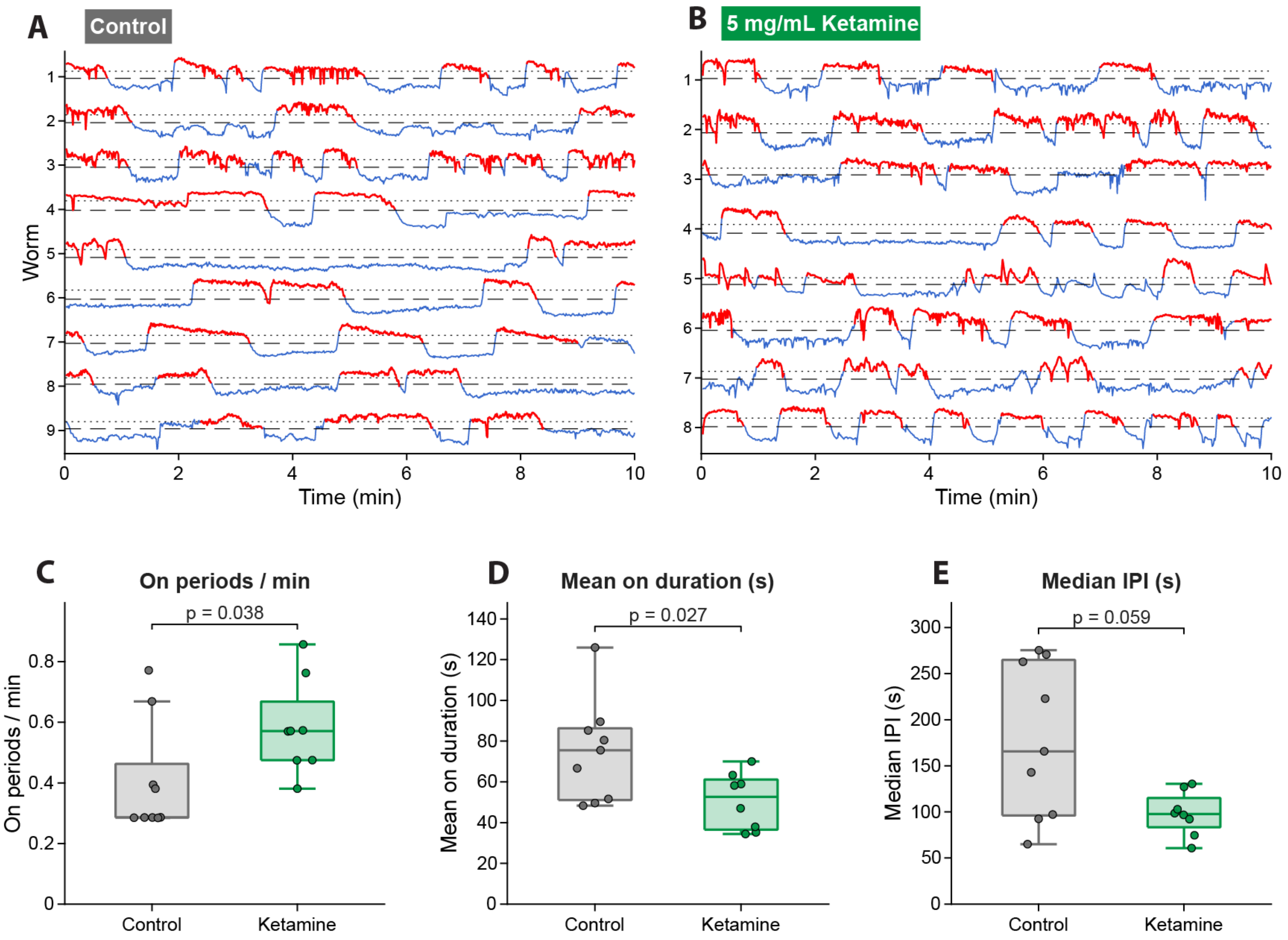

### Supplemental Figure 2

**Supplemental Figure 2**

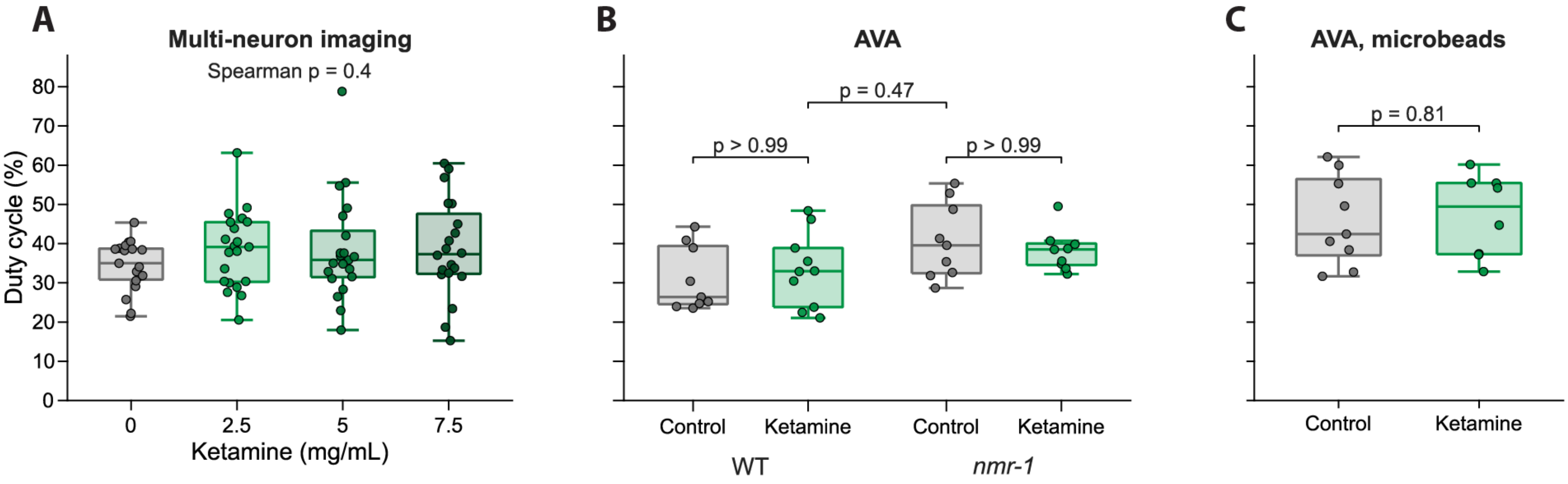
